# Graphene oxide/silver nanoparticle ink formulations rapidly inhibit influenza A virus and OC43 coronavirus infection *in vitro*

**DOI:** 10.1101/2021.02.25.432893

**Authors:** Meredith J. Crane, Stephen Devine, Amanda M. Jamieson

## Abstract

Respiratory tract infections present a significant risk to the human population, both through seasonal circulation and novel introductions with pandemic potential. There is a strong need for antiviral compounds with broad antimicrobial activity that can be coated onto filtration systems and personal protective equipment to augment their ability to remove infectious particles from the environment. Graphene oxide and silver nanoparticles are both materials with documented antimicrobial properties. Here, we tested the *in vitro* antiviral properties of several graphene oxide–silver nanoparticle composite materials, which were prepared through three different methods: reduction with silver salt, direct addition of silver nanospheres, and direct addition of silver nanospheres to thiolized graphene. These materials were tested over short time scales for their antiviral activity against two enveloped RNA viruses, influenza A virus and OC43 coronavirus, by performing viral plaque assays after exposure of the viruses to each material. It was found that the graphene oxide – silver nanoparticle materials generated by direct addition of the silver nanospheres were able to completely inhibit plaque formation by both viruses within one minute of exposure. Materials generated by the other two methods had varying levels of efficacy against influenza A virus. These studies indicate that graphene oxide-silver nanoparticle composite materials can rapidly neutralize RNA viruses and demonstrate their potential for use in a wide range of applications.

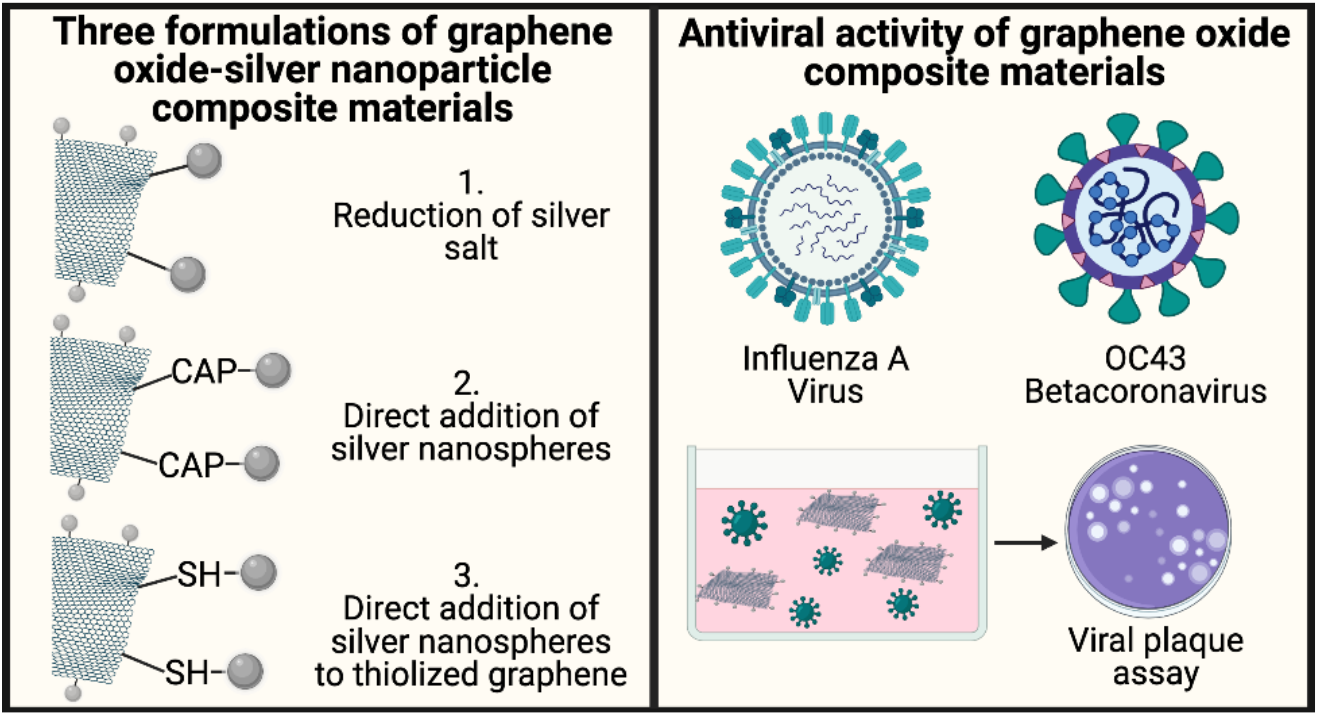

## Introduction

Viruses that infect the respiratory tract are a leading cause of morbidity and mortality in the human population^1^. These viruses range from those that circulate seasonally, such as the seasonal influenza viruses, and those that represent novel introductions into the population with pandemic potential, as experienced with the novel 2009 H1N1 influenza A virus lineage and the coronavirus SARS-CoV-2, the viral agent of COVID-19 disease^2–5^. While there is much focus on and an urgent need for the development of targeted therapeutics and vaccines for these illnesses, there is equal necessity to develop more novel, broad-spectrum antiviral compounds^6–12^. Such materials could be used to decontaminate surfaces or augment air filtration systems and personal protective equipment (PPE). For example, while certain masks, such as N95 masks, are protective through their ability to filter out a significant portion of harmful bacteria and viruses,^13,14^ few can actually kill viruses^15^. Similarly, gloves prevent contact of viruses with the skin, but do not prevent the virus from being passed to other surfaces. Supplementing these systems with broadly acting antiviral materials would compound their efficacy in protecting against infection.

Metal nanoparticles, such as silver, have well recognized antimicrobial properties. Silver nanoparticles (AgNPs) have been studied primarily for their antimicrobial potential against bacteria but have also proven to be active against several types of viruses including human immunodeficiency virus, hepatitis B virus, herpes simplex virus, respiratory syncytial virus, and monkeypox virus. The antiviral effect of silver nanoparticles for some viruses have been reported previously^16,17^. Recently, it has been suggested that AgNPs bind with the external membrane of lipid-enveloped viruses to prevent infection. Others have shown that silver nanoparticles are more effective as an anti-viral than silver ions alone^18,19^.

Graphene oxide (GO) is also a documented antimicrobial material. Research studies have shown that the two-dimensional structures, sharp edges, and negatively charged surfaces of GO can kill bacteria and viruses by disruption and/or oxidation of the plasma membrane^20,21^. In addition, the high surface areas of graphene and graphene oxide enable them to be loaded with high levels of antiviral agents, making them ideal drug carriers. Crucially, the combination of graphene or GO and antiviral agents has been shown to both increase their antiviral performance and reduce their toxicity, enabling significantly higher antiviral performance^22^.

Multiple reports have demonstrated antiviral and antibacterial properties of GO modified by the addition of AgNPs (GO-AgNP)^21,23–25^. One report, using porcine reproductive and respiratory syndrome virus, revealed that the antiviral activity mechanism of GO-Ag nanocomposites worked by preventing viral entry and by activation of the antiviral innate immune response. Furthermore, the GO-AgNP nanocomposite was more efficacious than either GO or AgNPs alone^24^. This synergistic antiviral effect of GO-AgNPs was also demonstrated against both enveloped and non-enveloped viruses by Chen *et al*^26^.

The work described here examined the antiviral properties of GO-AgNP composite materials developed by Graphene Composites as part of their patent-pending GC Ink antiviral formulations. A variety of materials were generated using three different production methods: reduction with silver salt, direct addition of Ag nanospheres, and direct addition of Ag nanospheres to thiolized graphene. These materials were tested for their ability to reduce the infectivity of two RNA viruses, influenza A virus (IAV) and the human coronavirus (HCoV) OC43, after short exposure periods. IAV is an enveloped virus of the orthomyxovirus family with a segmented single-stranded RNA genome ^27^. HCoV-OC43 is an enveloped betacoronavirus with a single-stranded RNA genome associated with the common cold in humans ^28^. It was found that many of the materials rapidly reduced infection levels, and that the materials produced by direct addition of Ag nanospheres completely inhibited the infectivity of both viruses. Together, these data reveal the potential for GO-AgNP composite materials to function as antiviral nanosystems that may enhance current infection prevention measures and antiviral therapies.

## Materials and Methods

### Preparation of graphene oxide

All production methods and GO-AgNP composite materials described in the Materials & Methods are covered by Graphene Composites’ patents pending. GO (1wt% in water, William Blyth) was used as received and shaken before use to homogenize the slurry. All final ink dispersions had the target concentration of 1mg/ml, so the graphene oxide was diluted 1/10 by mass with deionized water, to a nominal concentration of 0.1 wt%. Probe sonication in an ice bath fully dispersed and exfoliated the graphene oxide before decoration with AgNPs.

### Generation of silver nanoparticle-modified graphene oxide by reduction of silver salt

AgNPs were produced *in situ* by the reduction of Ag salts. Silver nitrate (99% purity, Alfa) was used as the silver salt and ascorbic acid (99% purity, Alfa) was used as the reducing agent. After preparation of the GO solution as described above, the pH was adjusted with either sodium hydroxide solution (0.1M) or ammonium hydroxide (1M), to pH 11, giving the optimum control of the particle size. The pH and temperature of the solution, as well as the addition rate of the reducing agent dictates the size of AgNPs produced. The reduction was carried out using the reducing agent in a 1 to 10 mole equivalent of the silver nitrate. Three different levels of silver decoration were achieved with nominal 20, 40, and 60 wt% Ag on GO levels through varying the amount of silver nitrate used in the reaction (materials R1, R2, and R3, respectively, Table 1). The resultant material was vacuum filtered and washed several times to remove excess salts and free silver particles.

**Table 1:** Composition of GO-Ag NP Materials.

| SAMPLE | FORMULATION |
| --- | --- |
| R1 | 90°C reduction of silver nitrate at pH 11 with ascorbic acid (20wt% Ag) |
| R2 | 90°C reduction of silver nitrate at pH 11 with ascorbic acid (40wt% Ag) |
| R3 | 90°C reduction of silver nitrate at pH 11 with ascorbic acid (60wt% Ag) |
| D1 | Direct addition of 10nm Ag NPs with capping agent to GO (2.5wt% Ag solid) |
| D2 | Direct addition of 20nm Ag NPs with capping agent to GO (2.5wt% Ag solid) |
| T1 | 10nm silver NPs on thiolized GO (4% silver) |
| T2 | 40nm silver NPs on thiolized GO (4% silver) |

### Generation of silver nanoparticle-modified graphene oxide by direct addition of silver nanospheres

10nm functionalized (capped) AgNPs were commercially sourced (Sigma-Aldrich). The chemical capping agent is adsorbed on the surface of AgNPs formerly on the coordinate bond. This effect provides the space steric hindrance and stabilizes the colloid system. 10nm or 20nm capped AgNPs (materials D1 and D2, respectively, Table 1) at a concentration of 0.02 mg/mL were directly added to GO to produce Ag-decorated GO via hydrogen bonding that occurs between the capping agent and the oxygen-based chemistry on the surface of the GO. The D1 material was manufactured at a nominal silver content of 2.5 wt% and D2 at 55 wt%. The manufacturing of this decorated material was undertaken using ultrasonic processing which enables both thorough mixing and exfoliation of the graphene oxide material to single or few layers as well as supplying sufficient energy to encourage the hydrogen bonding phase of the reaction to occur. The process resulted in a very stable dispersion being formed with no evidence of unbound silver particles.

### Generation of silver nanoparticle-modified graphene oxide by direct addition of silver nanospheres to thiolized graphene

Graphene oxide was chemically thiol functionalized at O, OH, and COOH sites using sodium hydrosulfide. The resultant thiolized GO particles were then filtered and washed to remove unreacted sodium hydrosulphide, then redispersed with water and mixed with commercially sourced AgNPs (Sigma Aldrich), which coordinate to the thiol groups via hydrogen bonding. 10nm and 40nm AgNPs were sourced (0.02mg/ml concentration Sigma Aldrich) and are designated as materials T1 and T2, respectively (Table 1).

### Thermogravimetric analysis of silver nanoparticle-modified graphene oxide

The 1mg/ml GO-AgNP solutions were pre-dried to 2-6% solid before analysis. The residual water was then driven off by initially heating to 120°C, then heated to 900°C to burn off the graphene leaving the residual silver. The silver content is determined as:

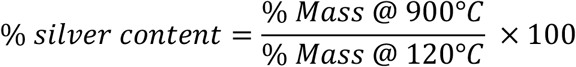

### GO-AgNP treatment of IAV

Use of IAV is covered under the Brown University Biohazards Research Authorization Protocol 20190802-032. GO-AgNP solutions (Table 1) were sonicated for 20 minutes to disperse the particles. The GO-AgNP solutions, vehicle, or 1x PBS were then added to a 96-well plate in a volume of 250μL. For treatments, GO-AgNPs were tested undiluted (100% GO-AgNP; H_2_O vehicle) or diluted 100-fold in 1x PBS (1% GO-AgNP; 1% H_2_O vehicle).

10^4^ or 10^5^ PFU of influenza A virus (A/WSN/33 (H1N1), IAV) was added to treatment wells in a volume of 50μL, for a final volume ratio of GO-AgNP:IAV of 5:1. 1x PBS supplemented with calcium and magnesium (Genessee Scientific) containing 0.2% BSA (w/v, Fisher Scientific) vehicle was added to control wells. 1, 5, or 10 minutes after the addition of IAV or vehicle, the plate was centrifuged at 1650xg for 5 minutes to remove GO-AgNPs. The supernatant was transferred to a clean 96-well plate. Each treatment condition was tested in triplicate.

### Quantification of IAV infectivity

Plaque assays were performed to measure IAV infectivity. MDCK cells were grown to 90-95% confluence in 6-well tissue culture plates in DMEM (ThermoFisher Scientific/Gibco) containing 10% FBS (ThermoFisher Scientific/Gibco) and 1% Penicillin-Streptomycin (ThermoFisher Scientific). The supernatant from the GO-AgNP treatment plates was serially diluted in 1x PBS supplemented with calcium and magnesium (Genesee Scientific) containing 0.2% BSA (Fisher Scientific) and 100μL was seeded onto MDCK cell monolayers. Inoculated plates were incubated for 1 hour at 37°C. Following the incubation, the inoculum was replaced with 1x DMEM containing 1.2% NaHCO_3_, 0.2% BSA, and 1% bacteriological agar (Oxoid). After 72 hours of incubation at 37°C, the 1x DMEM/1% bacteriological agar mixture was removed and the MDCK cell monolayers were stained with 0.1% crystal violet. The number of plaques per well was counted to determine viral titers.

### GO-AgNP treatment of human coronavirus OC43

Use of HCoV-OC43 is covered under the Brown University Biohazards Research Authorization Protocol 20190802-032. Treatments of human coronavirus (HCoV)-OC43 with material D2 were performed exactly as described for IAV except that the inoculating dose of HCoV-OC43 was 10^5^ PFU.

### Quantification of HCoV-OC43 infectivity

Vero E6 cells (ATCC CDL-1586) were grown to 80% confluence in 6-well tissue culture plates in DMEM (ThermoFisher Scientific/Gibco) containing 5% FBS (ThermoFisher Scientific/Gibco) and 1% Penicillin-Streptomycin (ThermoFisher Scientific). Supernatant from the HCoV-OC43 treatment plates was serially diluted in 1x PBS supplemented with calcium and magnesium (Genesee Scientific) containing 0.2% BSA (Fisher Scientific). 100μL of diluted virus was seeded onto Vero E6 cell monolayers and incubated for 1 hour at 33°C. The inoculum was then removed and 2mL of 1x DMEM containing 1.2% NaHCO_3_, 0.2% BSA, and 1% bacteriological agar (Oxoid) was added to each well. Plaque assays were incubated at 33°C for 5 days, after which the agar mixture was removed, and the cell monolayers were stained with 0.1% crystal violet. The number of plaques were counted to determine viral titers.

### Assessment of MDCK cell viability

MCDK cell viability following exposure to GO-AgNP supernatants or vehicle was assessed using a Cytotoxicity Detection Kit (Millipore Sigma #11644793001). Prior to testing, GO-AgNP supernatants or vehicle samples were serially diluted in 1x PBS/0.2% BSA exactly as done in preparation for plaque assays. 100μL of the serially diluted GO-AgNP supernatants or vehicle was seeded onto MDCK cells grown to 90-95% confluence in 6-well tissue culture plates. Plates were incubated for 1 hour at 37°C. Supernatants were then replaced with phenol red-free culture media. Culture media samples were obtained at 1- and 20-hours post-treatment and tested for the presence of lactate dehydrogenase (LDH) according to the manufacturer’s instructions. Briefly, an equal volume of cell culture supernatant was mixed with diaphorase/NAD^+^ catalyst containing iodotetrazolium chloride and sodium lactate dye solution in the wells of an optically clear 96 well plate. The mixture was then incubated for 30 minutes at 25°C. Cells grown only in the presence of media provided a low LDH control. Cells lysed with 2% Triton-X provided a high LDH control. Optical density (OD) was measured at 490nm with a reference wavelength of 650nm. The percent cell viability was calculated as:

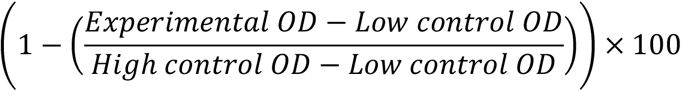

## Results

### Generation of AgNP-modified GO by reduction of silver salt

Scanning electron microscopy was used to determine the particle size of the Ag nanoparticles decorating the surface of the GO in materials R1, R2 and R3 (Table 1). The morphology and decoration pattern (for 60% silver) is shown in Figure 1A. The particle size of the Ag increased as the relative weight loading percentage increases (Figure 1B). The particle size distributions of the three materials also demonstrated that increasing the level of silver created a loss of control of the particle size of the Ag particles being produced, increasing from 53nm (mean) to 104nm (mean) over the declared range (Figure 1B).

**Figure 1.**
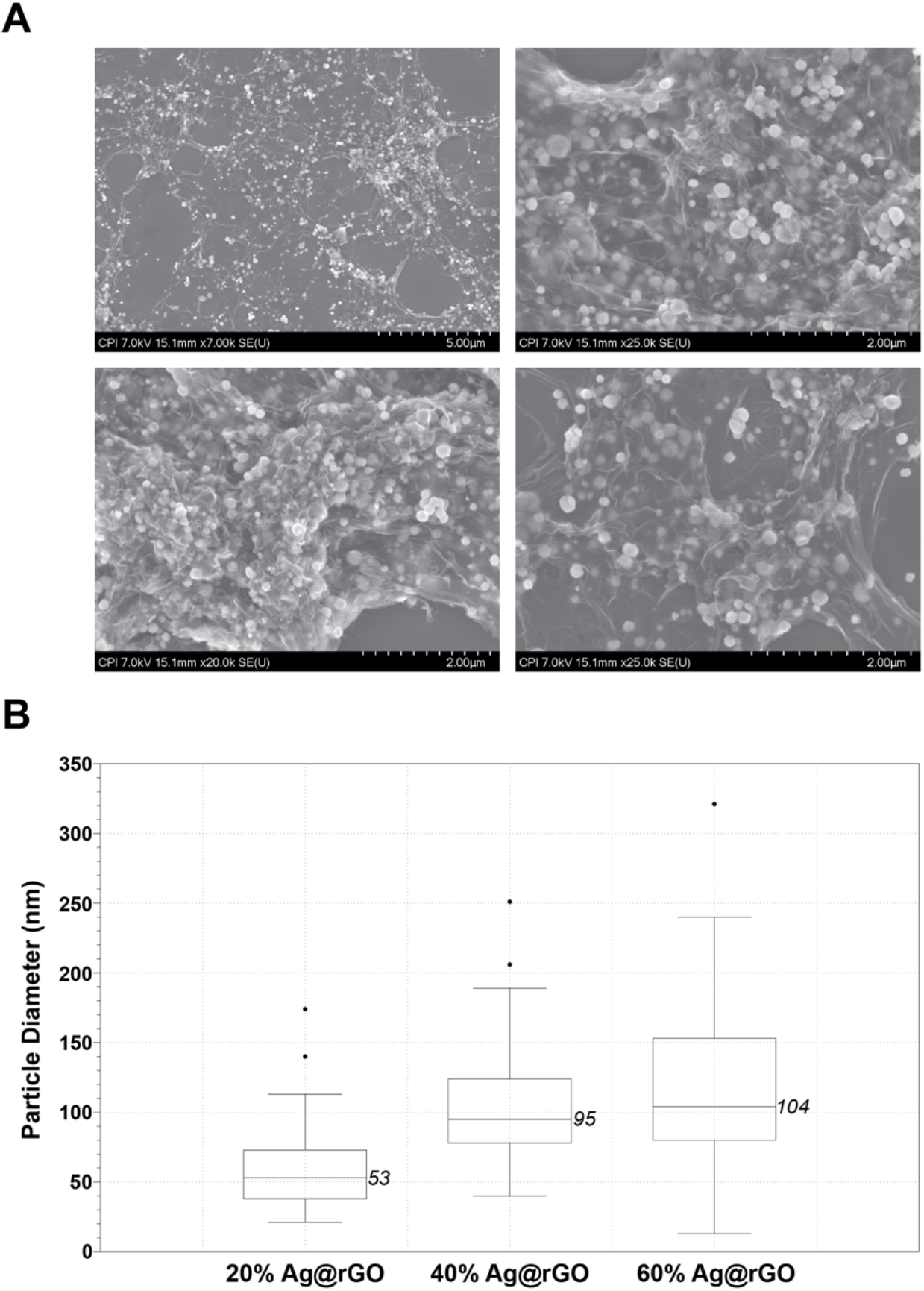
Generation of GO-AgNP by reduction of silver salt. (A) SEM micrographs of material R3 (60% Ag). (B) Ag particle size distribution of materials R1 (20%), R2 (40%), and R3 (60%).

The silver content of materials R1, R2, and R3 was determined by thermogravimetric analysis (Table 2). The actual silver contents observed were, in all cases, higher than the nominal amounts calculated (45%, 25%, and 10% higher for R1, R2 and R3 respectively). This is likely to be due to dispersion variation in the graphene oxide wt% levels of the William Blyth material at 1 wt% which are then exaggerated by diluting to 0.1 wt% or measurement error due to small procedural sample sizes.

**Table 2:** Calculated Ag Content.

| SAMPLE | Nominal % Ag Content | Dispersion % Ag Content | % Ag Content |
| --- | --- | --- | --- |
| R1 (20% Ag on GO) | 20 | 3 | 29 |
| R2 (40% Ag on GO) | 40 | 7 | 50 |
| R3 (60% Ag on GO) | 60 | 11 | 66 |

### Generation of AgNP-modified GO by direct addition of silver nanospheres

The Ag-decorated GO surface of material D1 was visualized by SEM (Figure 2A). The majority of Ag decoration on this material occurred around the periphery of the GO plates, which is where the majority of the oxygen containing chemistry occurs. The silver particles were well distributed across the dried surface examined in the SEM. Measurement of Ag particle size revealed no significant difference between the size distribution before and after processing (within measuring error) indicating that there had been no chemical change in the particles (Figure 2B). Thermogravimetric analysis showed the silver content to be 47% of a 0.3% solids content in the dispersion.

**Figure 2.**
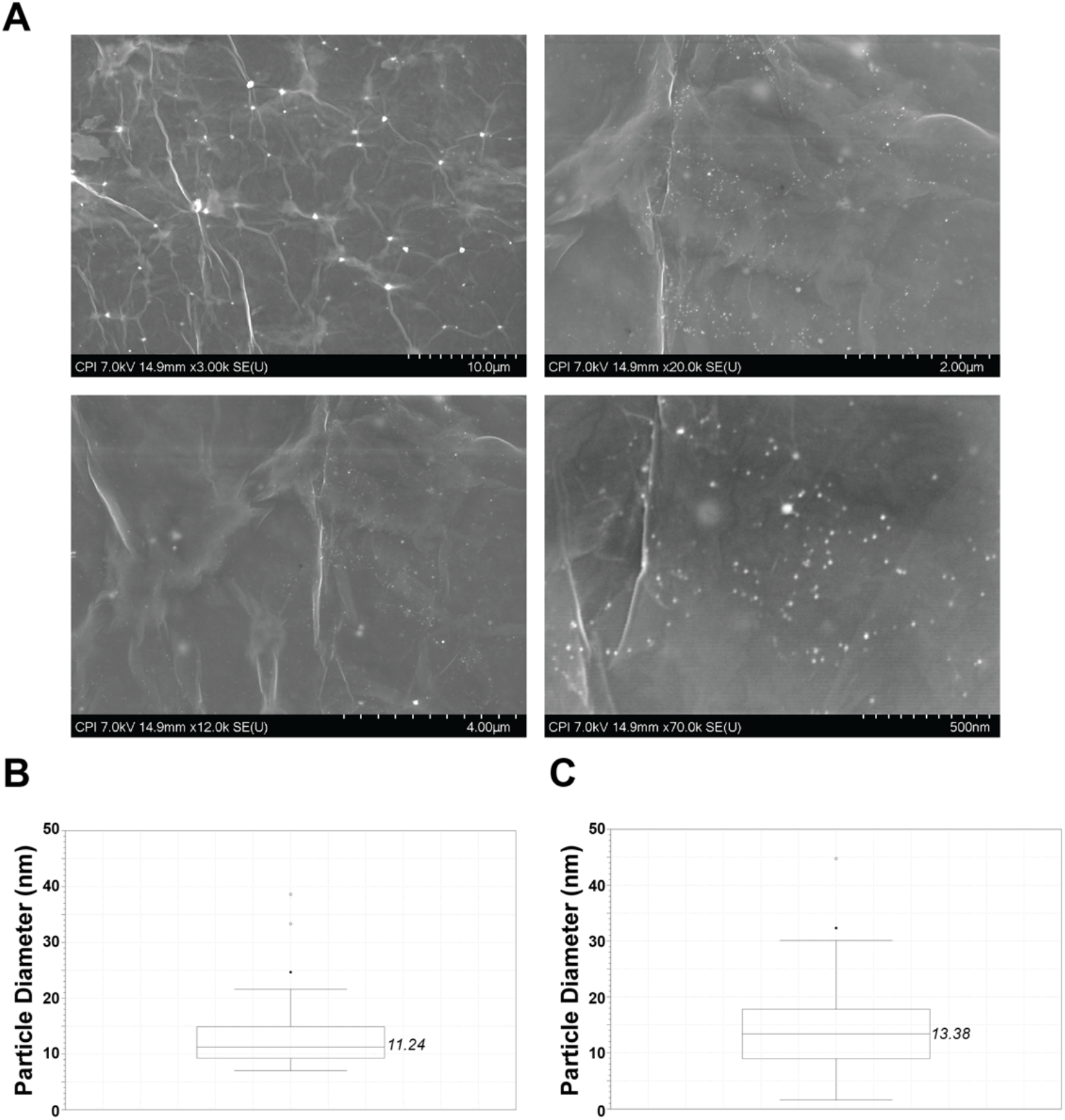
Generation of GO-AgNP by direct addition of Ag nanopsheres. (A) SEM micrographs of material D1. (B) Ag particle size distribution of material D1 before processing and (C) after processing.

### Generation of AgNP-modified GO by direct addition of silver nanospheres to thiolized graphene

The morphology and Ag decoration characteristics for materials T1 (10nm Ag) and T2 (40nm Ag) were examined by SEM analysis (Figure 3A and 3C). The particle size of the Ag particles decorating the GO surface was also measured for these materials (Figure 3B and 3D). In both cases, no noticeable change in the size was observed when compared to the materials as received. There was also evidence of agglomeration of Ag particles on the surface of the GO from both 10nm (T1) and 40nm (T2) materials.

**Figure 3.**
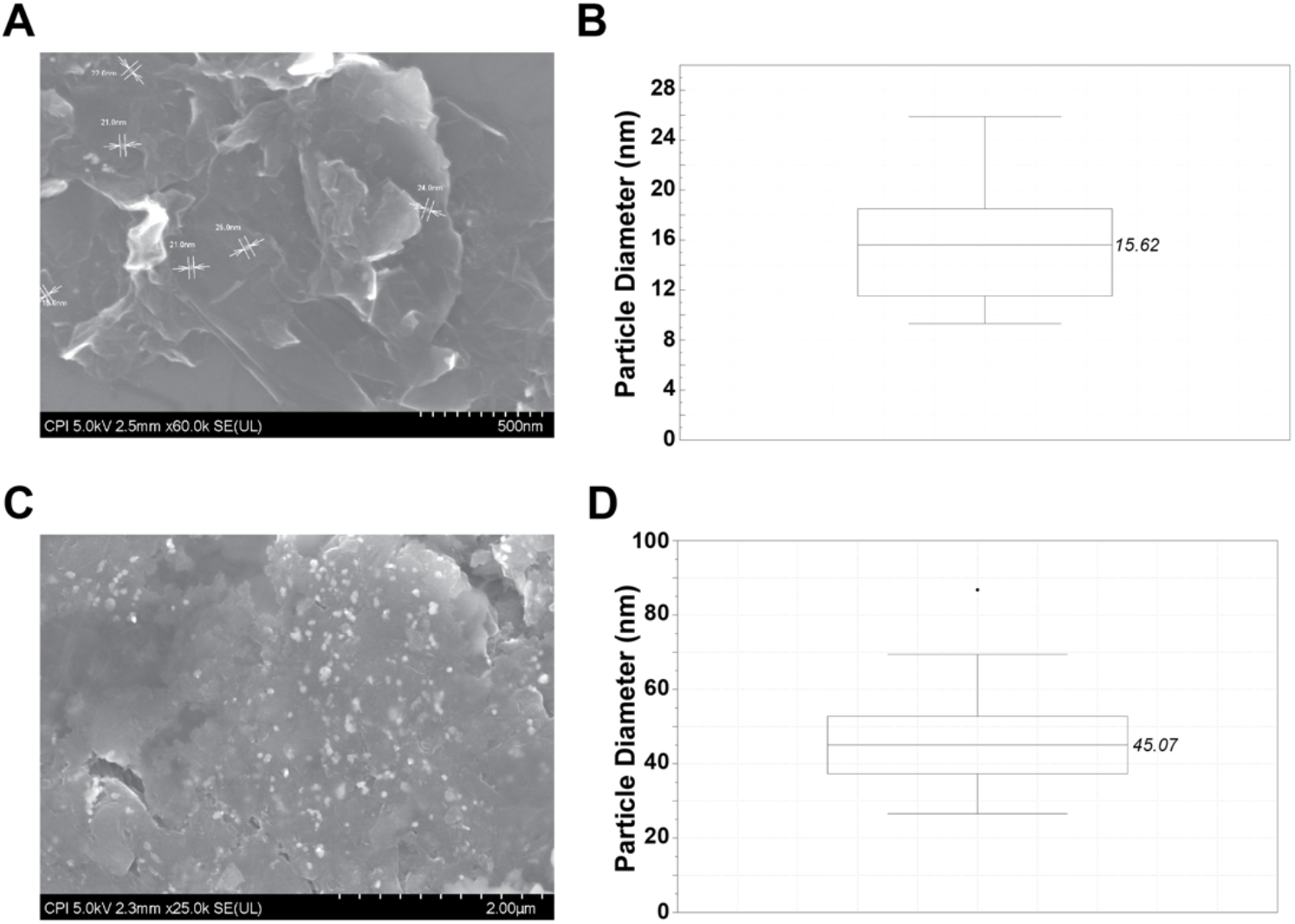
Generation of GO-AgNP by direct addition of Ag nanopsheres to thiolized graphene. (A) SEM micrograph and (B) Ag particle size data of material T1. (C) SEM micrograph and (D) Ag particle size data for sample T2.

### Effects of GO-AgNP materials on IAV infectivity

The seven GO-AgNP solutions with differing compositions as listed in Table 1 were tested for their ability to reduce the level of infectious IAV over time. GO-AgNP solutions were mixed with either 10^4^ or 10^5^ PFU IAV at a volume ratio of 5:1 and incubated for 1, 5, or 10 minutes. The solutions were tested in their undiluted state (100%) or were diluted to a 1% solution to determine the range of their potency. Plaque assays were then performed to assess IAV infectivity following GO-AgNP exposure. Two of the three materials prepared by the reduction of silver salt (materials R1 and R3) demonstrated a modest ability to inhibit IAV plaque formation by up to 0.5 log under certain treatment conditions compared to PBS alone. In contrast, material R2 did not alter plaque formation compared to PBS-treated groups (Figure 4A, B, and C).

**Figure 4.**
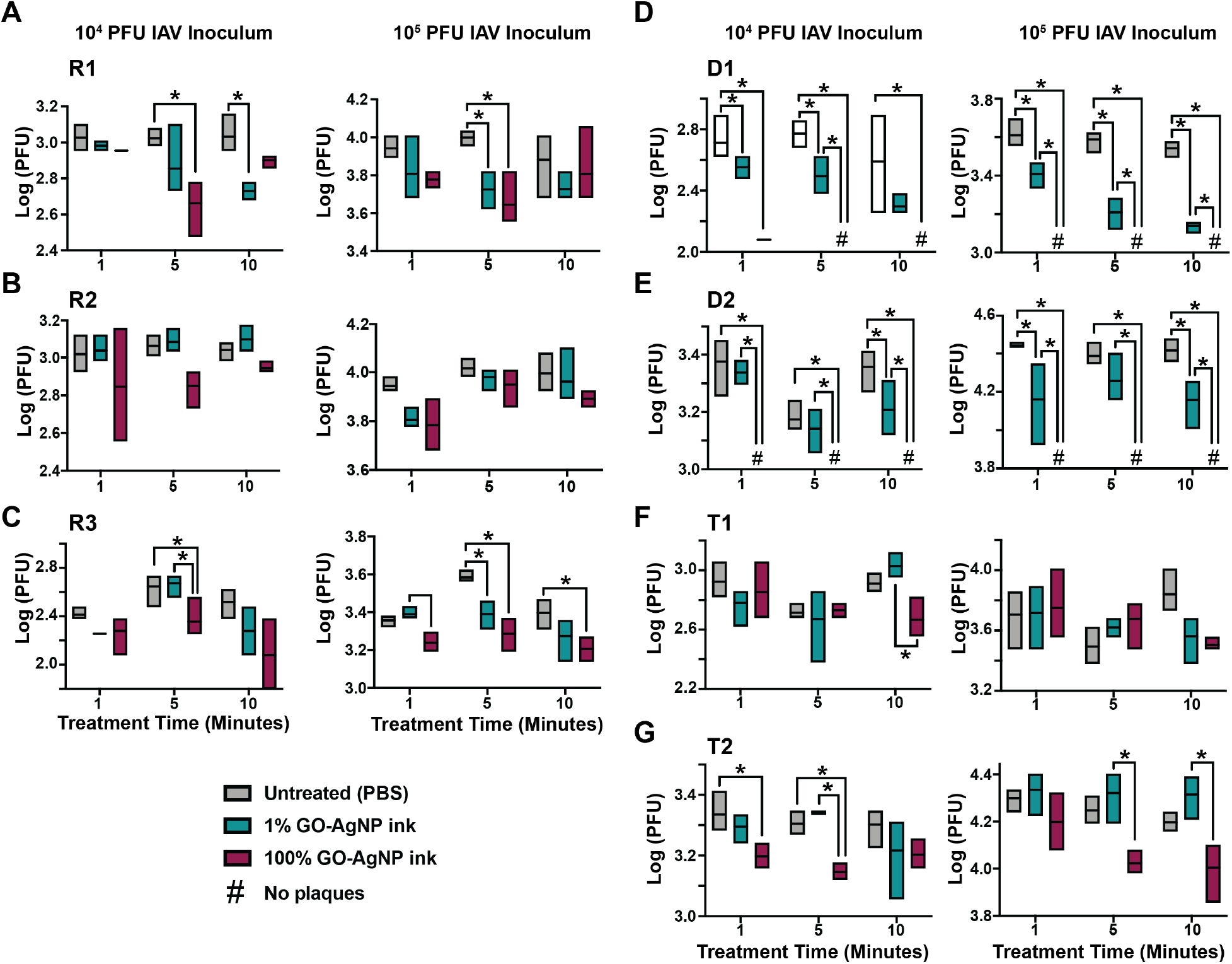
GO-AgNP exposure reduces IAV infectivity. IAV (10^4^ or 10^5^ PFU) was mixed with 1% (teal bar) or 100% (maroon bar) GO-AgNP solutions or 1x PBS (grey bar) for 1, 5 or 10 minutes. GO-AgNPs were then removed by centrifugation. Plaque assays were performed using MDCK cells to quantify viral plaque forming units (PFU) after treatment. Plaque assay results are shown following treatment with (A) sample R1, (B) sample R2, (C) sample R3, (D) sample D1, (E) sample D2, (F) sample T1, and (G) sample T2. Data shown are the average ± SD, n = 3 samples per group. * indicates p ≤ 0.05. Significance was determined using a two-way ANOVA with Tukey’s multiple comparison’s test.

The materials prepared by the direct addition of silver nanospheres to GO demonstrated a strong ability to reduce IAV plaque formation. Undiluted (100%) material D1 completely inhibited IAV plaque formation at all exposure time points examined. Exposure to a 1% solution of material D1 also significantly inhibited plaque formation at all time points examined (Figure 4D). Similarly, no IAV plaques formed after treatment with undiluted (100%) sample D2 at all time points examined. A 1% solution of sample D2 also reduced IAV plaque formation after 10 minutes of co-incubation (Figure 4E).

The two materials prepared by the direct addition of silver nanospheres to thiolized graphene (T1 and T2) inhibited IAV plaque formation up to 0.5 log under certain treatment conditions. Only the 100% solutions of these materials were successful in reducing viral plaque formation (Figure 4F and 4G).

Effects on virus viability were not due to the 1% or 100% GO-AgNP aqueous vehicles, as treatment with the vehicles alone did not alter IAV infectivity after 10 minutes of exposure (Supplemental Figure 1). To confirm that MDCK cell viability was not affected by exposure to GO-AgNP supernatants during the plaque assay, cell monolayers were exposed to the supernatants of each GO-AgNP material for 1 hour. MDCK cell viability following treatment was assessed by the release of LDH. GO-AgNP supernatants did not alter LDH release by MDCK cells 1 or 20 hours after exposure (Supplemental Figure 2).

### Effect of GO-AgNP material D2 on HCoV-OC43 infectivity

Tests with IAV revealed that materials D1 and D2 had the highest antiviral activity. Therefore, material D1 was further tested for antiviral efficacy against HCoV-OC43. One- and ten-minute exposure of HCoV-OC43 to 100% solutions of material D1 almost completely or completely prevented plaque formation (Figure 5), similar to what was seen with IAV. Ten-minute exposure to a 1% solution also had a significant antiviral impact (Figure 5B).

**Figure 5.**
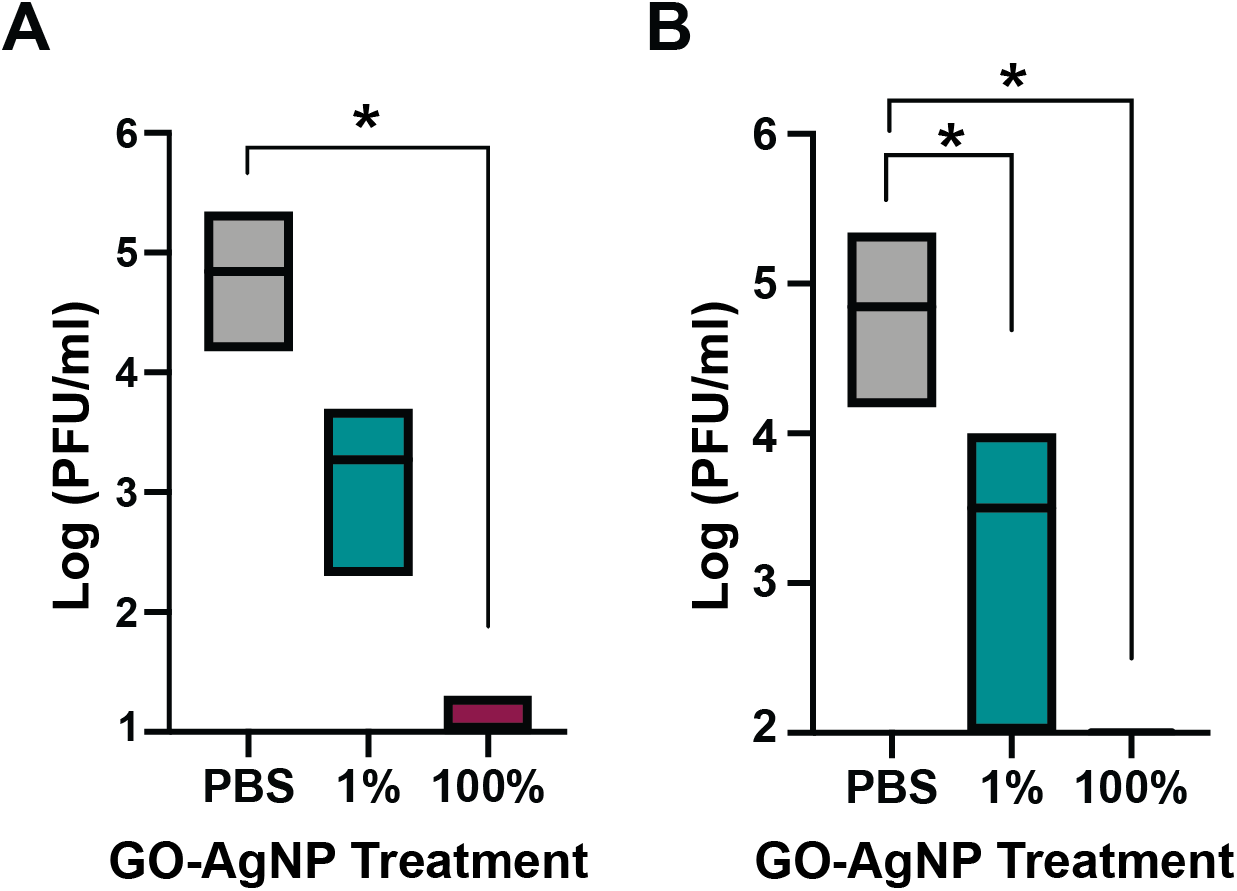
GO-AgNP exposure reduces HCoV-OC43 infectivity. HCoV-OC43 was treated with PBS vehicle (grey bar), or a 1% (teal bar) or 100% (maroon bar) solution of material D1 for 1 minute (A) or 10 minutes (B). Plaque assays were performed using Vero E6 cells to quantify viral plaque forming units (PFU) following treatments. Data shown are the average ± SEM, n = 9 replicates per group from 3 biological replicates. * indicates p ≤ 0.05. Significance was determined using a two-way ANOVA with Tukey’s multiple comparison’s test.

## Discussion

The studies described here identify two GO-AgNP materials with potent and rapid antiviral activity in solution against two enveloped RNA viruses associate with human respiratory infection. A total of seven materials were tested, which resulted from three different production methods. The remaining five materials possessed a range of modest to no antiviral effects against IAV.

The differences in measured antiviral effect are consistent with chemical and morphological differences among the seven tested samples. One such difference involved the stabilization of the AgNPs. Materials D1 and D2, which exhibited the greatest antiviral effect, were created by the direct addition of commercially sourced Ag (capped) nanospheres. The non-reactionary chemistry during the manufacture of these materials leaves the surface chemistry of the graphene oxide intact. This allows for maximum potency of the material in terms of the reduction of viral load ^29^. In addition, this capping agency presence has two functions; namely, it allows steric hindrance stabilization of the colloid as the capping agent adsorbs onto the coordinate bond of the silver, and it allows hydrogen bonding of the capping agent to the oxygenated species on the surface of the GO. Samples T1 and T2 similarly used commercially sourced AgNPs, but in contrast, these materials had no surface functionalization or capping agents present so had a tendency to aggregate during the coordination with the thiol group phase of reaction.

The reaction state of the graphene oxide also differed among the three production methods. In all but samples D1 and D2, the GO was reduced to reduced GO (rGO). Samples R1, R2 and R3 had a marginal reduction of the GO to rGO during the reduction phase, which reduces silver nitrate to silver. This is in agreement with Li *et al* ^29^, who showed that the oxidation state and reactive surface groups on the planar surface dominate the toxicity to lipid membranes, with pristine GO performing considerably better than rGO. Similarly, with samples T1 and T2, the GO is reduced to rGO during the thiolization process. This, as postulated by Liu^21^, would reduce the ability of the GO to oxidize the plasma membrane of the viral particles. Lastly, the role of the ascorbic acid in the reduction reaction is critical, as it controls the size of the AgNPs produced. A faster addition of the reducing agent and speed of reaction results in smaller particles being produced; thus, the resultant size of the Ag nanoparticles produced is proportional to the level of Ag required, i.e. the reaction takes longer for the higher loading levels allowing more time for the growth of particles.

As reported in previous studies, the rapid and high antiviral activity of materials prepared by the direct addition of AgNPs to GO likely results from a synergistic effect between the GO and the AgNPs. This is consistent with the slower reaction times for the materials containing rGO, as the AgNPs would be the only component of these materials undertaking oxidation of the lipid membrane. Importantly, materials D1 and D2 were able to fully inhibit plaque formation for two different human-tropic RNA viruses after just one to five minutes of treatment prior to the infection stage of the assay. Rapid disruption of the viral envelope is one likely mechanism of action, as prior studies have shown the ability of GO materials to disrupt membranes through physical and/or oxidative stress^20,21,29^; this suggests that these materials would also be effective against other enveloped viruses. When considering the use of these materials in coatings for PPE, high-touch surfaces, and air filtration units, rapid virus destruction is a critical factor, as demonstrated here. Taken together, this work demonstrates the potential use of GO-AgNP materials in a variety of contexts to reduce human exposure to dangerous viral pathogens.

## Conclusion

GO-AgNP composite materials that were prepared by the direct addition of silver nanospheres (capped) to GO were found to prevent *in vitro* infection with IAV virus and HCoV-OC43 after 1 to 10 minutes of exposure in solution. These findings demonstrate the ability of these materials to rapidly reduce the infectivity of two respiratory viruses, highlighting their potential use as antimicrobial agents in a variety of applications.

## Ancillary Information

### Supporting Information

Supplemental Figure 1: GO-AgNP aqueous vehicles do not alter IAV infectivity.

Supplemental Figure 2: GO-AgNP supernatants do not affect MDCK cell viability.

### Author Contributions

MJC and AMJ designed and performed infection experiments and wrote the manuscript. SD provided experimental materials and materials characterization data.

## Acknowledgements

The authors would like to thank Sandy Chen for helpful discussion and insight regarding experimental design and reporting of results and Dr. Adriana Kajon for providing the OC43 coronavirus strain. The abstract graphic was created using BioRender.com. Funding for this work was provided by the Brown University COVID-19 Seed Fund. No financial support was provided by Graphene Composites, Ltd.

## Abbreviations Used

AgNP: silver nanoparticle
GO: graphene oxide
rGO: reduced graphene oxide
IAV: Influenza A virus
HCoV: human coronavirus
MDCK: Manin-Darby canine kidney
PPE: personal protective equipment
LDH: lactate dehydrogenase

