## Supporting Information for "Graphene oxide/silver nanoparticle ink formulations rapidly inhibit influenza A virus and OC43 coronavirus infection *in vitro*"

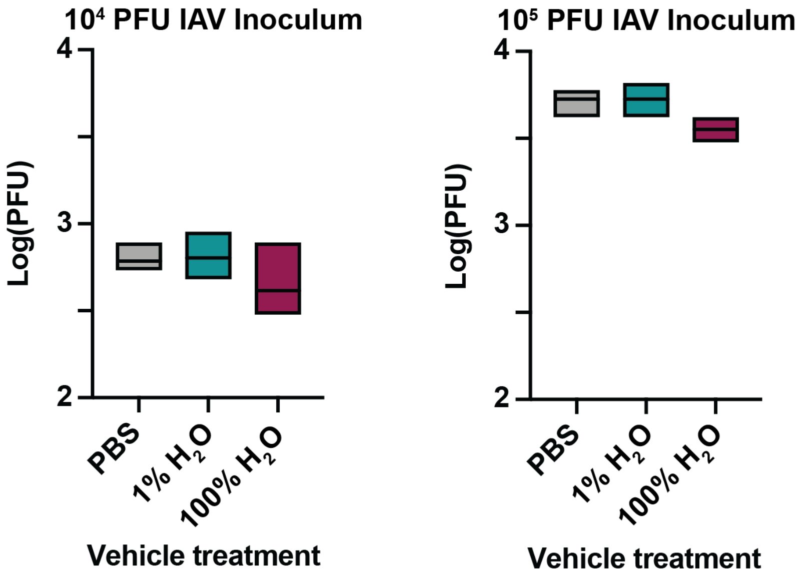


**Supplemental Figure 1.** GO-AgNP aqueous vehicles do not alter IAV infectivity. $10^4$ (left) or $10^5$ PFU IAV (right) was exposed to 1% $H_2O$ (diluted in 1x PBS, teal bar) or 100% $H_2O$ (maroon bar) vehicle or to 1x PBS (grey bar) for 10 minutes. Viral PFUs were then measured by plaque assay. Data shown are the average ± SD, n = 3 samples per group. Significance was determined by a Kruskal-Wallis test.

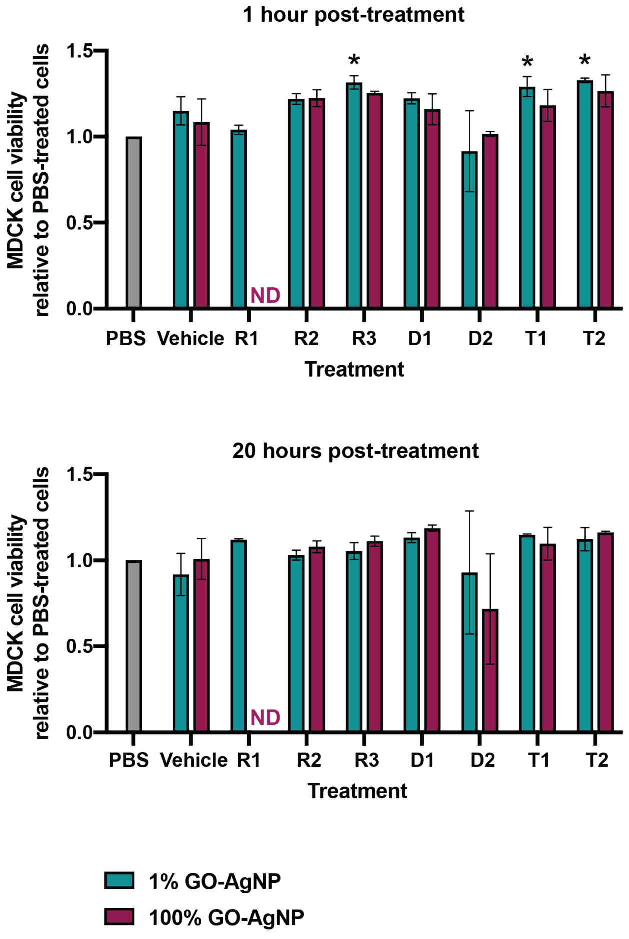


**Supplemental Figure 2.** GO-AgNP supernatants do not affect MDCK cell viability. MDCK cells were exposed to the supernatants of each GO-AgNP sample, vehicle, or 1x PBS (grey bar). Teal bars show MDCK cells treated with the supernatant from a 1% solution of GO-AgNPs or 1% $H_2O$ vehicle, while maroon bars indicate 100% GO-AgNPs or 100% $H_2O$ vehicle. The concentration of LDH was measured in media collected 1 hour (top) and 20 hours (bottom) post-treatment. The percent viability was determined as described in the Materials and Methods and was normalized against the PBS control sample. Data shown are the average ± SD, n = 3 samples per group. * indicates $p \leq 0.05$ as compared to the PBS control. Significance was determined using a two-way ANOVA with Tukey's multiple comparison's test. ND = no data available.
